# Representational drift reflects ongoing balancing of stochastic changes by Hebbian learning

**DOI:** 10.1101/2025.01.05.631363

**Authors:** Jens-Bastian Eppler, Thomas Lai, Dominik Aschauer, Simon Rumpel, Matthias Kaschube

**Affiliations:** Frankfurt Institute for Advanced Studies, Frankfurt, Germany; Institute of Computer Science, Goethe University, Frankfurt, Germany; Institute of Physiology, Focus Program Translational Neurosciences, University Medical Center, Johannes Gutenberg University Mainz, Mainz, Germany; Centre de Recerca Matematica, Barcelona, Spain

## Abstract

Recent evidence indicates that even under stable environmental and behavioural conditions responses to sensory stimuli undergo continuous reformatting over the course of days, a condition described as representational drift. However, the processes underlying this phenomenon remain poorly understood. Examining the dynamics of signal and noise correlations among neuron pairs in chronic calcium imaging experiments in the mouse auditory cortex, we investigate how activity-dependent, Hebbian-like plasticity, and activity-independent, stochastic synaptic processes contribute to representational drift. We found that signal correlations predict future noise correlations, suggesting that stimulus-induced co-activation leads to increased effective connectivity between neuron pairs. Moreover, simple linear network models were able to account for the observed temporal dependencies between signal and noise correlations, but only if both Hebbian-like plasticity and stochastic changes of either inputs or recurrent synapses contribute to representational drift. In conclusion, our findings suggest that continuous sensory input-driven Hebbian-like plasticity can balance ongoing stochastic synaptic changes, thereby preventing the network’s functional degradation.

## Introduction

Although the same sensory stimulus is generally thought to evoke a consistent and stable percept over time, studies show that the corresponding neuronal activity patterns can undergo substantial changes over the course of days, a phenomenon known as representational drift. In recent years evidence for representational drift has been observed in various areas of the brain [1], [2], [3], [4], [5] suggesting to reflect a fundamental property of neuronal circuits with possible implication for various cognitive functions [6]. Despite its prevalence, the processes underlying representational drift remain largely unresolved.

Synaptic changes in network connectivity are a likely driver of changes in stimulus-evoked activity patterns [7]. Traditionally, synaptic changes have been linked to activity-dependent Hebbian processes, where specific patterns of pre-and postsynaptic activity can modify the strength of synaptic connections between neurons. This form of plasticity is crucial for learning and memory formation [8], and recent modelling studies have suggested that Hebbian learning may also contribute to representational drift. Ongoing Hebbian plasticity may continuously modify the current network configuration [9], which could cause sensory representations to drift over time. This concept is akin to statistical learning, where organisms continuously adapt and potentially overwrite previously established memories with new information from constantly changing inputs [10]. Other models explain drift by ongoing Hebbian learning in a degenerate solution space during noisy representation learning [11], [12]. These modelling studies are in line with recent experimental findings, which suggest that not only experience, but also time can influence the nature and degree of representational drift [13], [14], [15].

However, synaptic connections also exhibit substantial and seemingly stochastic fluctuations independent of neuronal activity, observed both vitro and in vivo [16], [17]. These ongoing changes in synapses were also observed under conditions where neuronal activity was silenced [18], [19], [20] possibly reflecting fluctuations in the dynamic molecular makeup of individual synapses [21]. Thus, alongside Hebbian synaptic plasticity, activity-independent, stochastic changes may also influence the stability of activity patterns [7], although their role has been less studied to date. Furthermore, the relationship between these two forms of synaptic changes and their respective contributions to representational drift remain unclear.

Signal and noise correlations can provide a sensitive readout to changes in the activity and structure of a network. Foundational studies have explored the roles of synchronous activity and neuronal correlation in neural binding and visual processing [22], [23], as well as in the reliability and efficiency of neural coding [24], [25], [26], [27]. Many studies distinguish between signal correlations, which reflect the tuning similarity of neuron pairs [28], [29], and noise correlations, which capture the shared variability in neuronal responses independent of external stimuli. Studies on the origins of noise correlations suggest that they can depend on mono-synaptic and multi-synaptic connections, recurrent pathways, and stimulus independent shared inputs [24], [30], [31] suggesting their interpretation as effective connectivity between neurons (for a comprehensive review of both signal and noise correlations see [27]). Despite their importance and broad applications, the long-term dynamics of both signal and noise correlations over days remain largely unexplored.

Importantly, signal and noise correlations may be affected differently by Hebbian-type, experience-dependent plasticity than by stochastic, experience-independent synaptic changes. Large signal correlations could via Hebbian plasticity promote the establishment of stronger synaptic connections between correlated neurons, and thus predict changes towards enhanced noise correlations between such neurons. Conversely, stochastic synaptic drift may also affect both signal and noise correlations, but without any systematic temporal relationship between the two. Thus, changes in signal and noise correlations could potentially reveal important constraints on the plasticity processes underlying representational drift, but so far this possibility has been explored very little.

In this study, we utilize signal and noise correlation to shed light on the relative contributions of the different forms of synaptic changes to representational drift. We analysed chronic calcium imaging data of sound evoked responses in the mouse auditory cortex [5], a model well suited to address this question as both synapses [16], [32], [33], [34] and neuronal representations [5], [35], [36], [37] have been documented to undergo changes under stable environmental and behavioural conditions. We found that signal correlations predict future noise correlations, but not vice versa. The simplest network model that could account for these findings combined a Hebbian-like learning process in recurrent synaptic connections with a component of stochastic changes: either from drifting inputs or stochastic changes within the recurrent connections themselves. Furthermore, when examining the dynamics of signal and noise correlations during auditory cued fear conditioning, we observed a diminished influence of signal correlations on noise correlations. Together, these results suggest that Hebbian learning counterbalances ongoing stochastic changes during basal conditions, whereas the impact of stimulus driven changes is slightly reduced after fear conditioning.

## Results

To test the role of synaptic changes underlying representational drift, we analysed our publicly available dataset [38] obtained using chronic two-photon calcium imaging in 80 fields of view in layers 2/3 of 10 awake, head-fixed mice. Neurons were expressing the calcium indicator GCaMP6 together with a structural nuclear marker enabling high-fidelity tracking of individual neurons across several days. Animals were passively listening to 20 trials of 34 short stimuli (50 to 70 ms) in randomized order in four imaging sessions separated by two days each. Cells were tracked across all imaging sessions and neuronal recordings were pre-processed resulting in activity traces for each of the 18,249 neurons (for a detailed description of recording procedures and data handling see [5] and Methods). Between the second and third imaging session (i.e. days 3 and 5), the mice underwent a fear conditioning paradigm on day 4, where one of the stimuli was paired with a mild electric shock (Fig. 1a). An example field of view is shown in Fig. 1b. Neuronal somata are labelled in red and GCaMP6m expression is shown in green. Example activity traces for two cells can be seen in Fig. 1c, both single trial activity in black and median activity across trials in purple (for further examples see Fig. S1).

**Figure 1:**
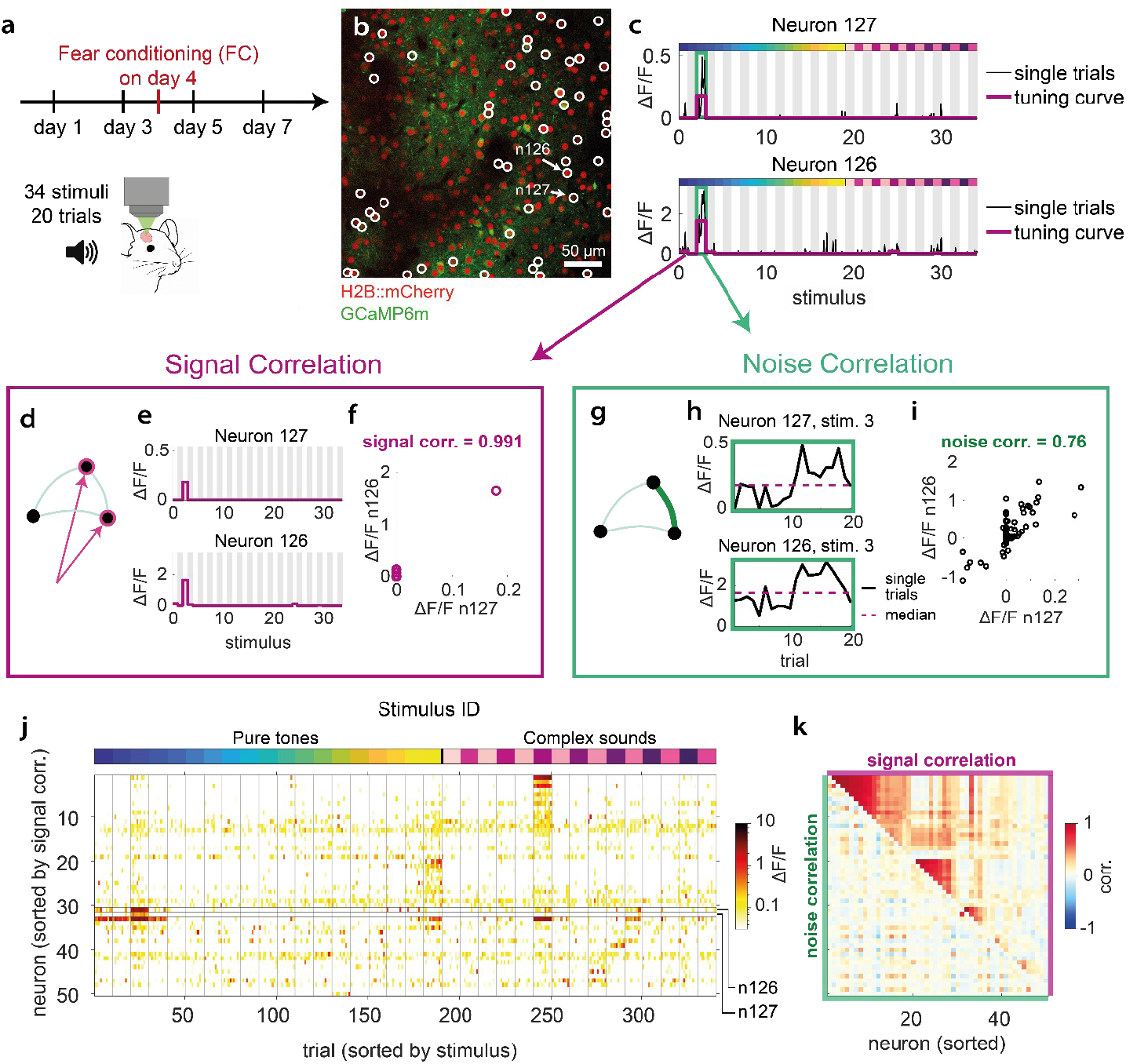
Experimental data and estimation of signal – and noise correlations. a) Experimental timeline. b) In vivo two-photon image of a local population of neurons in layer 2/3 of the auditory cortex showing expression of GCaMP6m (green) and H2B::mCherry (red) (for details see [5]). Arrows indicate location of the neuron pair shown in (c), (e), (f), (h) and (i). White circles show location of the neurons shown in (j) and (k). Neurons were selected for high signal correlation, noise correlation or activity. c) Single trial activity (black line) and median stimulus-specific response over trials (purple line) of the example neuron pair with particularly high signal and noise correlation shown in (b). Green box indicates the stimulus responses displayed in (h). Grey and white columns indicate stimuli. d) Sketch illustrating signal correlations as a measure of tuning similarity. e) Tuning curves of the example neuron pair in (c). f) Tuning curve of neuron 126 vs. neuron 127. g) Sketch of the interpretation of the noise correlation as a measure of an effective connectivity between neurons. h) Single trial activity of the example neuron pair in (c) in response to stimulus 3 (Zoom-in on the green boxes shown in (c)). Purple line represents median response over trials. i) Trial-to-trial variability for all trials and stimuli of neuron 126 vs. neuron 127. j) Activity traces of 50 example neurons which have a maximum stimulus response of >0.4 and a maximum signal- or noise correlation of >0.75 (marked by white circles in (b)). Neurons are sorted by hierarchically clustering the signal correlation matrix, trials are sorted by stimuli (stimulus ID is indicated by the colorbar above). Stimulus set consisted of 19 pure tones from low (blue) to high (yellow) frequency and 15 complex sounds (purple). k) Signal- and noise correlation matrix of the neurons shown in (j). Lower triangular matrix shows pairwise noise correlations, upper triangular matrix shows pairwise signal correlations.

We analyzed signal correlations, reflecting the stimulus-driven input to the cortical network (Fig. 1d-f) and noise correlations which served as a proxy for effective cortical interactions (Fig. 1g-i). Signal correlations between pairs of neurons were obtained by first computing each neuron’s median response across trials for each stimulus (Fig. 1e) and then calculating the Pearson correlation coefficient between tuning curves for each neuron pair (Fig. 1f). To compute noise correlations, we first subtracted for each stimulus the median activity from each single trial response (Fig. 1h). Computing the Pearson correlation coefficients of the residual activity between all pairs of neurons then yielded the noise correlations (Fig. 1i). To obtain a more robust estimate of both signal and noise correlations, we cross-validated our estimates via bootstrapping (see Methods for details). Fig. 1j shows for a representative subset of cells from an example field of view their stimulus-evoked activity and Fig. 1k their corresponding pairwise signal and noise correlations. Note that high signal correlations between pairs of neurons at a given imaging time point do not necessarily indicate high noise correlations and vice versa.

### A dynamic equilibrium of signal and noise correlations during representational drift

To explore the dynamics of signal and noise correlations, we first plotted their distributions at a given day, and observed that both display a mode near zero and a tail extending toward positive correlations (signal correlations: Fig. 2a, noise correlations: Fig. 2d), consistent with previous reports [39], [40]. Both distributions were remarkably stable across all four imaging time points. Next, we investigated whether correlations were stable at the level of individual neuron pairs. To assess the stability of signal correlations, we first divided the range of correlation values into eight bins and assigned neuron pairs into these bins based on their signal correlation on the first imaging day. We then plotted the signal correlation distribution for each group separately on the next imaging day (day 3), while maintaining the group assignment from day 1. This revealed that signal correlations varied substantially across days and for each group correlations tended towards the original overall distribution (Fig. 2b). Plotting the mean signal correlation for each group at both time points illustrates the drift of these means towards a common mean (Fig. 2c). Moreover, applying an analogous procedure to noise correlations revealed that that these show similarly strong changes as signal correlations (Fig. 2e,f). Note that these changes across days are unlikely to be caused by experimental noise, as we estimated the noise level through cross-validation (Methods). Taken together, both signal and noise correlations were highly volatile under basal conditions across a time span of two days, while their overall distributions were maintained.

**Figure 2:**
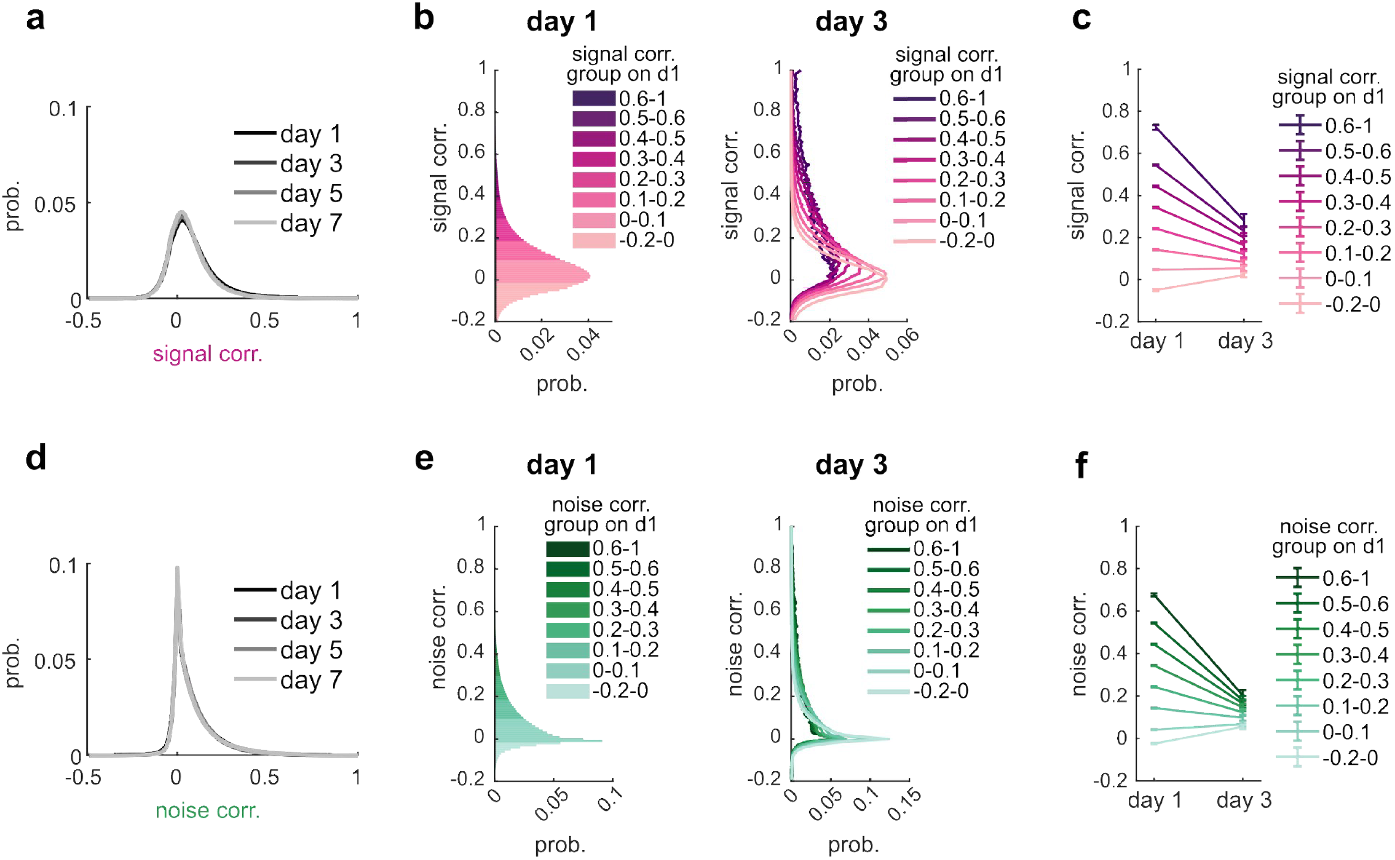
Signal and noise correlations are in a highly volatile state. a) Distribution of pairwise signal correlations across days. b) Left: Neuron pairs were assigned to groups based on their signal correlations on day 1. Color indicates threshold used for grouping (see legend). Probability normalized over all neuron pairs. Right: Signal correlation distribution of each group on day 3. Probability normalized over neuron pairs within groups. c) Mean signal correlation of each group on day 1 and day 3. Error bars represent SEM across neuron pairs. d) - f) Same as a) - c) but for noise correlations.

### Signal correlations predict future noise correlations

Having established that signal and noise correlations show substantial changes over time, we next wondered whether some of these changes were consistent with ongoing Hebbian learning. To address this question, we examined the relation between the changes in signal correlation and noise correlation. As Hebbian learning posits that two neurons strengthen their connectivity if they are co-activated, we hypothesized that neuron pairs co-activated by the same stimuli may exhibit stronger connectivity at a later time point due to Hebbian learning, which would tend to strengthen the noise correlation in such pairs. In this case, neuron pairs with high signal correlations should have increased noise correlations on the next imaging day.

To test this hypothesis, we investigated the extent to which the signal correlation levels of neuron pairs predicted their noise correlations on the next imaging day (Fig. 3a). To minimize the factor of noise correlation levels on day 1 in this analysis, we began by grouping neuron pairs based on their noise correlations, as we had done previously (Fig. 2f). In Fig. 3b we illustrate our analysis using the highest noise correlation group (NC = 0.6 to 1; Fig. 3b left). First, we subdivided this group into eight subgroups according to their signal correlation on day 1 (Fig. 3b right). Given the low correlation of signal and noise correlations on a given day (Fig. 1k, Fig. S3), neuron pairs in this group are expected to cover a wide range of signal correlations. For each signal correlation subgroup, we then computed the mean noise correlation on day 1 and on day 3 (i.e. the subsequent imaging day) and found that subgroups with high signal correlations on day 1 exhibited systematically higher noise correlations on day 3 than those with low signal correlations (Fig. 3b middle). At the same time, noise correlations differed only little on day 1, indicating that this effect cannot be explained by differences in noise correlation on day 1. We applied the same analysis to the other noise correlation groups and found that such temporal dependency between signal and noise correlations across days was also evident for the lower noise correlation groups albeit to a lesser degree (the three highest NC groups are depicted in Fig 3c; all groups are shown in Fig. S2a). These results indicate that cell pairs with a high noise correlation on day 1 tend to exhibit less decay in noise correlation from day 1 to day 3 if their signal correlation is high on day 1. Moreover, using the same analysis to assess the influence of noise correlations on future signal correlations revealed a weaker predictive effect (Fig. 3d, 3e). Although the subgroups with high noise correlation tended to have a slightly increased signal correlation on the next day, the differences in mean signal correlation across subgroups were low (Fig. 3f, for a complete overview of all subgroups see Fig. S2d).

**Figure 3:**
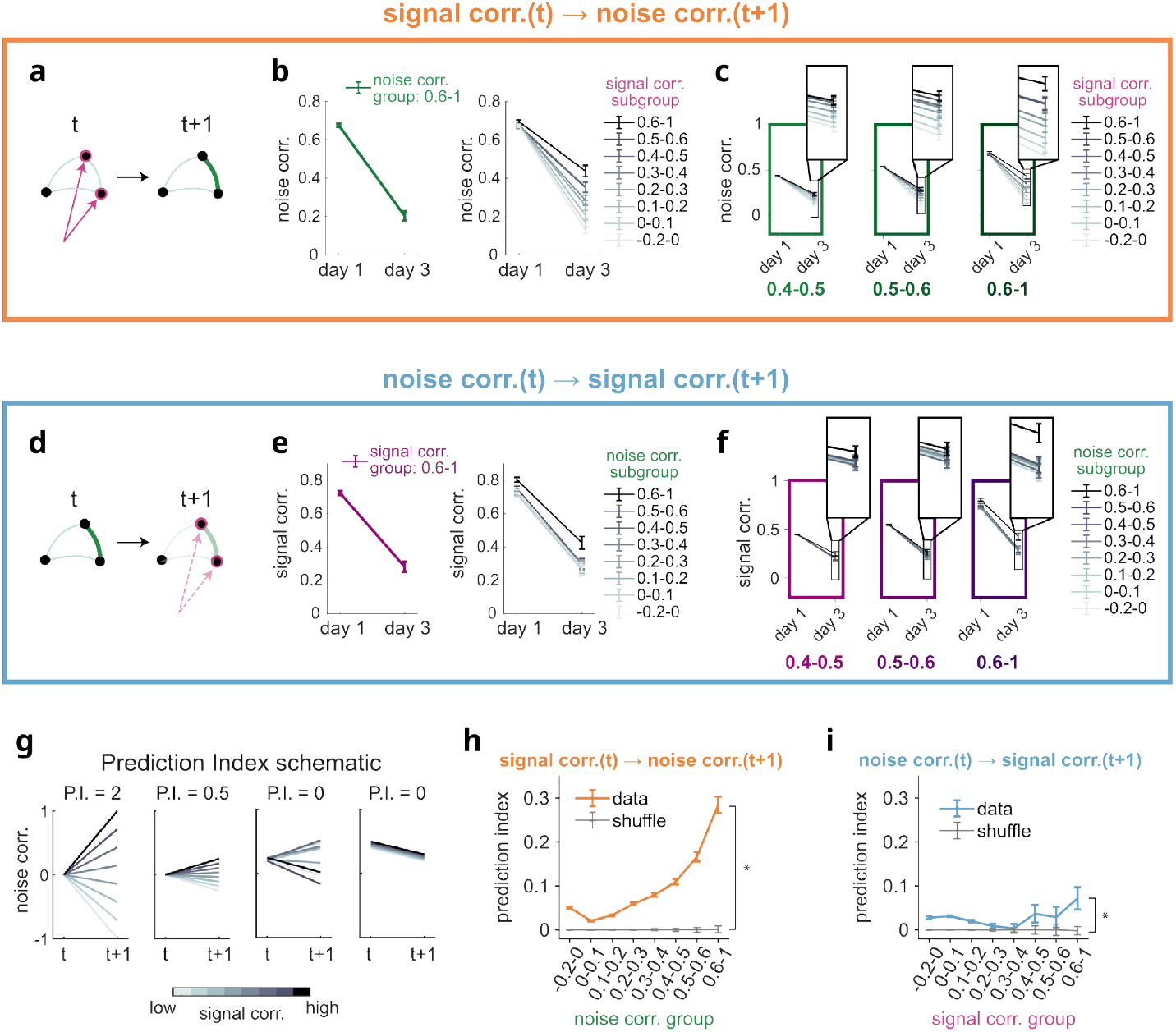
Signal correlations predict noise correlations under basal conditions. a)-c) Assessment of the predictive effect of signal correlations on noise correlations. a) Illustration of this effect. b) Left: mean noise correlation on day 1 and 3 of neuron pairs with noise correlation 0.6-1 on day 1. Right: mean noise correlations on day 1 and 3 of the same neuron pairs as on the left, but sub-grouped by their signal correlation. Individual lines represent individual signal correlation subgroups, colour indicates signal correlation on day 1. c) Same as in (b), but for two more noise correlation groups. Right subplot corresponds to the group shown in (b). All error bars in SEM. d)-f) Assessment of the predictive effect of noise correlations on signal correlations. Same as (a)-(c) but with reversed signal and noise correlations. g) Illustration of the prediction index (P.I.). It quantifies the predictive effect of signal correlations on noise correlations, or vice versa, by assessing the ordering and the spread of the lines due to the subgrouping. For illustration, the prediction index is indicated for four toy examples. h) Prediction index of signal correlation day 1 on noise correlation day 3 for the different subgroups defined in (b) (orange line, mean +-SD of bootstrapped distribution). Shuffle in gray (mean+-SD). i) Prediction index of noise correlation day 1 on signal correlation day 3 (blue line, mean +-SD of boostrapped distribution). Shuffle in gray (mean +-SD). *p<0.05 (WSR test).

To quantify these effects, we defined the prediction index (Fig. 3g; see Methods). The prediction index assumes a maximal value of 2 if the mean correlations on day 3 are maximally spread apart and sorted in the same order as the subgroups on day 1 (Fig. 3g, left, illustrates this for the case of noise correlations and subgroups based on signal correlations on the first day (coded by gray values). The prediction index reaches its minimal value of −2 if the spread is maximal in reverse order (Fig. 3g, middle left), while its value is 0 if there is no ordered spread (Fig. 3g, middle right) or no spread at all (Fig. 3g, right). Thus, the prediction index quantifies the extent to which the order of subgroups on day 1 corresponds to the spread of correlations on day 3, offering a metric for assessing the predictive power of signal correlations on future noise correlations and vice versa. Calculating this prediction index for each group, we found that under basal conditions, signal correlations have a significantly predictive effect on future noise correlations (Fig. 3h). The predictive effect of noise correlations on future signal correlations was significant, too, but substantially weaker than that of signal correlations on future noise correlations (Fig. 3i).

To corroborate these results with an alternative analysis method, we assessed the stabilizing relationship between signal and noise correlations by scattering signal (noise) correlations on day 3 against signal (noise) correlation on day 1, conditioned on noise (signal) correlations on day 1, and found that signal correlations contribute to the retention of noise correlations over time, while noise correlations have minimal influence on signal correlation stability (see Fig. S4 and S5). These results, thus, corroborate our findings in Fig. 3 and complement them by highlighting the stabilizing role of signal correlations on noise correlations.

Together, our empirical findings suggest that under basal conditions, signal correlations are predictive of future noise correlations, whereas noise correlations are barely predictive of future signal correlations. Such temporal relationship appears consistent with ongoing Hebbian plasticity induced by sensory input, which may thus be a strong factor underlying representational drift. To reveal its contribution, relative to other stochastic, input-independent factors, we study a network model in the following.

Notably, the observed temporal relationship was reduced when applying auditory cued fear conditioning (see Fig. S6), suggesting that this type of behavioural learning paradigm interferes with the ongoing remodelling of representations due to sensory-input driven Hebbian learning, possibly to facilitate the consolidation of learned associations into memory storage.

### Minimal circuit models suggest interplay of Hebbian and stochastic plasticity

To explore potential mechanisms underlying the observed dynamics in signal and noise correlation, including their asymmetric temporal relationship, we employed a linear recurrent network model (Fig. 4a). This model was designed to identify the minimal plasticity mechanisms that could explain our experimental results. The dynamics of the network are described by the following equation:

**Figure 4:**
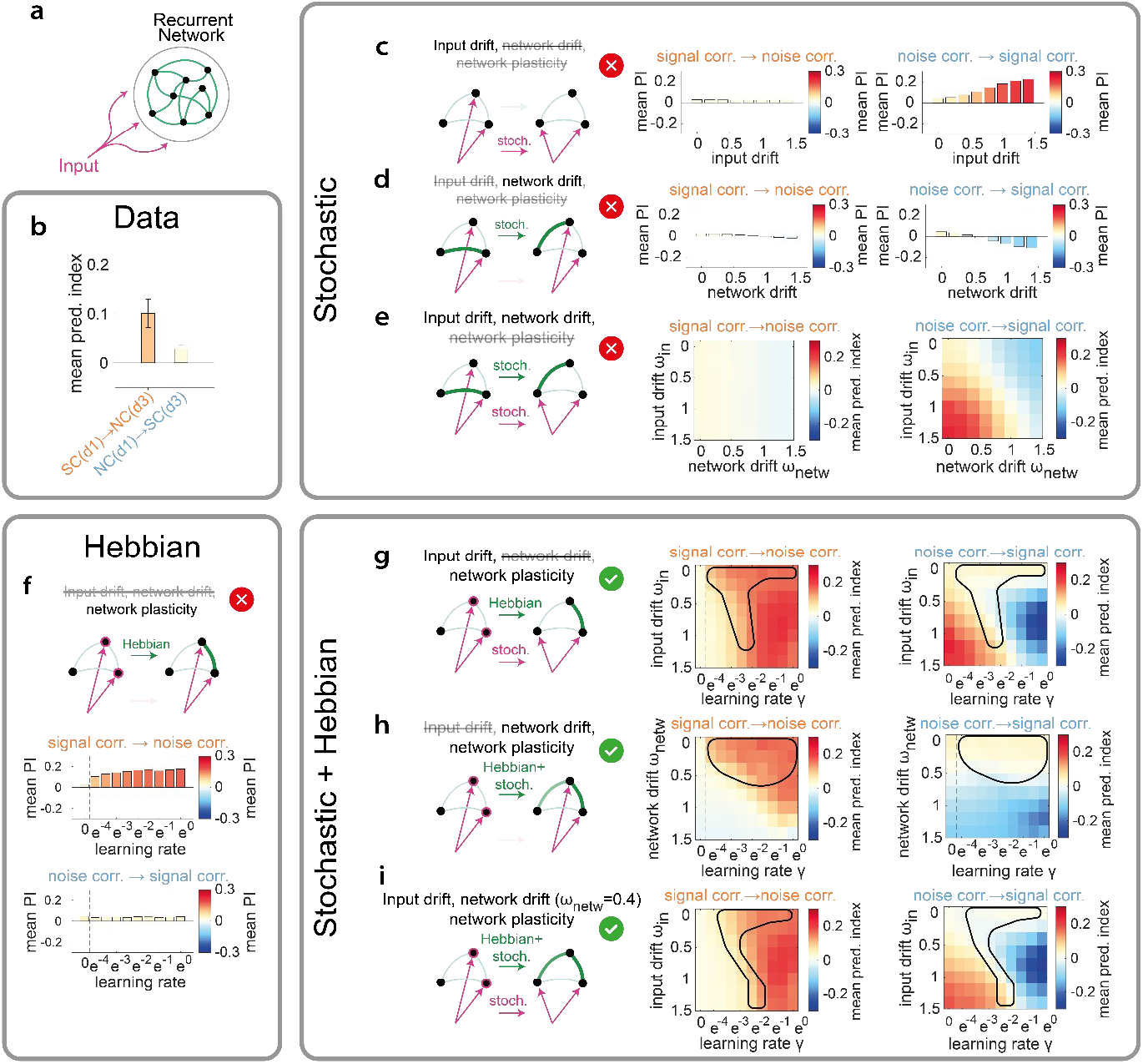
Circuit models suggest an interplay of Hebbian and stochastic plasticity is required to account for the experimental observations. a) Sketch of model architecture. b) Experimentally observed mean prediction index of signal correlations on noise correlations (left bar) and of noise correlations on signal correlations (right bar), summarizing Figure 3. Bar colour reflects mean prediction index using the same colour bar as in (c)-(i). c)-e) Models with stochastic changes. Mean prediction index in a model with stochastic input drift (c), stochastic network drift (d), and stochastic input and network drift (e). Left: sketch of network mechanism. Middle: mean prediction index of signal correlations on noise correlations. Right: mean prediction index of noise correlations on signal correlations. f) Model with Hebbian mechanism. Top: sketch of network mechanism. Middle: mean prediction index of signal correlations on noise correlations. Bottom: mean prediction index of noise correlations on signal correlations. g)-i) Model with stochastic and Hebbian mechanism. Mean prediction index in a model with stochastic input drift and network plasticity (g), stochastic drift and plasticity in the network weights (h), and all three mechanisms combined (i). Left: sketch of network mechanism. Middle: mean prediction index of signal correlations on noise correlations. Right: mean prediction index of noise correlations on signal correlations. Parameter scan in (i) shows results for one example network drift strength (*ω*_*net*_ =0.4). Black contours indicate plausible parameter spaces in agreement with data.

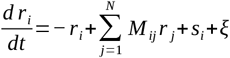

where *r*_*i*_ is the firing rate neuron *i, M*_*ij*_ represents the synaptic weight from neuron *j* to neuron *i, s*_*i*_ is the stimulus part of feedforward input to neuron *i*, ξ is additive Gaussian noise, and *N* is the total number of neurons in the network. To generate a dataset similar to experimental data we applied matching experimental parameters such that the same analysis procedures could be applied (see Methods, Fig. S8a-h).

In this model, we explored which plasticity mechanisms could account for the following key experimental observations: the high volatility of signal and noise correlations over time (Figs. 2b, 2c, 2e, 2f), the predictive effect of signal correlations on future noise correlations (Figs. 3h, 3i), and the stabilizing role of signal correlations on noise correlations (Figs. S4b, S4c). We assumed that synaptic weight changes in the network can occur through activity-dependent processes, such as Hebbian plasticity, or via stochastic activity-independent processes. We focused on three potential mechanisms, either in isolation or combined: (A) stochastic changes in the input, (B) stochastic changes in the recurrent network weights, and (C) input-dependent Hebbian-like plasticity in the recurrent connections.

We first tested whether stochastic changes alone could account for the observed dynamics of signal and noise correlations, considering changes only in the inputs, changes only in the recurrent connections, or a combination of both. If such stochastic changes alone were sufficient to account for our empirical observations, then this would suggest the possibility that these could be explained without the need for activity-dependent plasticity mechanisms like Hebbian learning. To test this hypothesis, we implemented stochastic changes for both inputs and recurrent connections in the model using an Ornstein-Uhlenbeck process [41] to introduce variability while maintaining overall statistics in a steady state. We varied the rate of change and computed the prediction index across subsequent time points (see Methods, Fig. S8i-n) to test for temporal dependencies between signal and noise correlation, and vice versa (Fig. 4). In addition, we performed the conditional correlation stability analysis (Fig. S10) analogously to the experimental data (Figs. S4, S5).

First, we examined the effect of stochastic changes in the input (Fig. 4c, left). This allowed signal correlations to fluctuate significantly, mirroring the volatility observed in experimental data. However, while the model could account for the changes in signal correlations, it failed to capture the changes in noise correlations (data not shown). Importantly, irrespective of the rate of synaptic changes, noise correlations depended little on signal correlations, with a prediction index close to zero (Fig. 4c middle), indicating that input drift alone cannot explain the observed interaction between signal and noise correlations. Next, we tested stochastic drift in the recurrent synaptic weights while keeping the input constant (Fig. 4d). Again, whereas such drift in the recurrent connections could account for the overall volatility (data not shown), the predictive effect of signal correlations on noise correlations remained absent (Fig. 4d middle). Finally, we combined stochastic changes in both the input and recurrent connections. As expected, this combination produced volatility in both signal and noise correlations, similar to what we observed experimentally, but still failed to reproduce the predictive effect of signal correlations on noise correlations, as the prediction index remained near zero (Fig. 4e middle). Moreover, for large combinations of drift rates, the model produced considerably positive or negative prediction indices for noise correlations on signal correlations (Fig. 4e right). These effects are primarily caused be strong recurrent connections, but are inconsistent with our experimental observations (Fig. 4b). Additionally, at higher levels of recurrent synaptic drift, we observed a reverse ordering of the correlations, likely due to the mean-reverting nature of the stochastic process, leading to extreme values being indicative of a smaller value at a consecutive time point and thus resulting in a negative prediction index (Fig. 4d). Thus, while stochastic processes, specifically in recurrent connections, are able to account for the high volatility of signal and noise correlations observed in the experimental data, they are insufficient to explain the predictive relationship between signal and noise correlations.

Given that stochastic processes alone could not explain the experimental data, we next implemented a simple Hebbian-like learning mechanism to investigate whether input-driven plasticity could reproduce the observed dynamics of signal and noise correlations. Hebbian-like learning was implemented as an interpolation between the connectivity matrix at the previous time step and a Hopfield term computed for the input vectors (see Methods). Using this Hebbian-like learning mechanism alone, the model was indeed able to reproduce the predictive effect of signal correlations on noise correlations (Fig. 4f). However, after several iterations, the network weights converged to the values imposed by the external input, causing both signal and noise correlations to stabilize and lose their volatility (data not shown).

Anticipating that stochastic changes could possibly counteract this restrictive tendency of a pure Hebbian-like learning mechanism [42], we next explored whether both processes together could explain the experimental results, by combining stochastic changes in the input with Hebbian-like plasticity in the recurrent network. In this model, the inputs were allowed to drift stochastically over time, while synaptic weights in the recurrent network were updated according to the Hebbian-like learning rule previously described (see Method for details). This combination successfully reproduced both key features: the ongoing volatility in signal and noise correlations (data not shown), as well as the predictive effect of signal correlations on noise correlations (Fig. 4g). The input drift introduced sufficient variability in the external stimuli, thus preventing the network from converging, while the Hebbian-like plasticity ensured that co-active neurons developed stronger connections, maintaining the predictive relationship between signal and noise correlations. We also examined a model combining stochastic changes in recurrent connections with Hebbian-like plasticity. We find that this combination also successfully reproduces both the volatility of signal and noise correlations and the predictive effect of signal correlations on noise correlations (Fig. 4h). In this model, the stochastic drift in the network drives continuous synaptic changes, while Hebbian-like plasticity prevents the network from becoming overly disorganized.

Furthermore, the combination of all three mechanisms, i.e. input drift, network drift, and Hebbian-like plasticity, also successfully reproduces the experimental observations (Fig. 4i). Similar findings were obtained using conditional correlation stabilization (see Fig. S4) instead of the prediction index (Fig. S10). In all three models, a strong correspondence with the experimental values was achieved for low drift rates in combination with intermediate learning rates.

In conclusion, a balance between stochastic changes (either in the feedforward inputs or the recurrent connections) and Hebbian-like learning in recurrent connections was required to reproduce our experimental findings in the model.

## Discussion

In this study, we investigated the mechanisms underlying representational drift in the mouse auditory cortex utilizing signal and noise correlations. Under stable environmental and behavioural conditions, we observed that both signal and noise correlations were highly volatile. Interestingly, signal correlations consistently predicted and stabilized noise correlations. Using a simple network model, we interpreted this pattern as Hebbian plasticity balancing an ongoing stochastic process in the inputs and/or the connections within a local network.

Signal and noise correlation have been studied extensively and shown to be an important characteristic of the functional properties of neural networks (e.g. [24], [30], [31]). Despite their importance in neural computation, relatively little is known about their long-term dynamics. Our findings showed that both signal and noise correlations are highly volatile under stable environmental and behavioural conditions. While this volatility might initially seem surprising, it is consistent with the well-documented perpetual remodelling of neuronal tuning properties termed representational drift [1], [2], [3], [4], [5]. This drift is likely resulting from changes in the underlying synaptic connectivity in the network [16], [17], [18], that can be observed even in the absence of neural activity [18], [19], suggesting it reflects an intrinsic property of synapses [20]. Importantly, the volatility in correlations we observe was not entirely random, but the predictive and stabilizing effect of signal correlations on noise correlations suggests the involvement of a Hebbian-like process in representational drift. In fact, our model shows that Hebbian plasticity in recurrent synaptic connections when combined with stochastic changes in connectivity can explain this stabilizing effect (Fig. 4g-i). This Hebbian plasticity can be thought of as stabilizing the network in the face of intrinsic fluctuations: In the presence of stochastic synaptic fluctuations, Hebbian mechanisms act to counterbalance this synaptic drift, with the input acting as a stabilizing force, preventing excessive disorganization that would result from stochastic changes alone. Such interpretation is in line with studies showing that Hebbian-like synaptic plasticity can stabilize network dynamics, ensuring functionally relevant connections to persist despite random synaptic turnover [11], [12], [43].

While such input-driven plasticity to counterbalance the effect of intrinsic stochastic synaptic drift appears plausible, it is important to note that part of the reorganization we observe could be induced by input-driven plasticity itself, given that the environment and the animal’s behaviour and brain state are constantly changing. In fact, even under the stable experimental conditions employed to obtain the data used in our study, the observed volatility in signal and noise correlations could in parts also be triggered by smaller changes in behaviour and external stimuli rather than by intrinsic stochastic processes only. This perspective is supported by previous studies emphasizing how networks update their representations as they encounter new inputs over time [9]. Despite the habituation of animals and the controlled conditions in our experiments, not only stochastic connectivity changes upstream of the auditory cortex could lead to changes in cortical inputs; also variations in sensory experience between imaging sessions could potentially shift input-driven synaptic changes over days. In our model, this latter scenario is also described by the case in which the inputs are assumed to change stochastically and Hebbian-like plasticity is applied to the recurrent network (Fig. 4h), but now with a different interpretation: Namely that these stochastic changes approximate the changing sensory inputs and that the network adapts to these new inputs updating its internal structure, reflecting a form of ongoing statistical learning. This is reminiscent of models proposed in language acquisition and sensory adaptation [44], [45], [46], [47]. This interpretation highlights the dynamic nature of cortical networks, where even in the absence of overt behavioural changes, the brain remains plastic, constantly integrating new information from its environment. Moreover, this interpretation suggests that representational drift is not merely a reflection of intrinsic fluctuations but is instead influenced by the constant influx of external stimuli. It underscores the idea that experience, rather than time alone, drives the continuous adaptation of neural representations, as supported by recent theoretical models and experimental findings [9], [13], [48].

Given that the balance of two antagonistic processes controls representational drift, we hypothesize that this balance may be shifted by internal and external processes leading to higher or lower drift rates. Indeed, increased exposure to external stimuli in an enriched environment can slow representational drift [4], [49]. This could be interpreted in a way that augmented reinforcement of consistent sensory inputs reduces the relative impact inherent stochastic synaptic changes. Future experiments isolating and manipulating individual components of inherent stochastic synaptic dynamics or forms of Hebbian plasticity are needed to determine how input-driven and network-driven processes interact to shape representational drift, shedding light on the mechanisms that support ongoing learning and adaptation.

The two suggested underlying causes of representational drift, input-driven changes and intrinsic synaptic processes, bear interesting links to mechanisms of forgetting. In fact, theories of forgetting, such as trace decay and interference theory, offer parallels to the two primary processes we observe. Trace decay suggests that memories fade over time due to intrinsic changes, consistent with synaptic drift [50], [51], [52], while interference theory posits that new information replaces old, aligning with the idea of continuous incorporation of new sensory inputs [53], [54]. A recent study in the mouse hippocampus supports both mechanisms, showing that drift in neuronal firing rates changes with time, while tuning properties of neurons change with experience [14]. The link between drift and forgetting has also been made in a recent review [55], and our findings are consistent with the notion that representational drift could be an ongoing process of both intrinsic decay and constant extrinsic overwriting, reflecting both aspects of forgetting.

Together, our findings highlight that data acquired from a mature and apparently stable brain on a given day, merely reflect a snapshot of two counterbalancing dynamic processes safeguarding functionality in an inherently unstable network.

## Methods

### Experimental Data

We analyzed data from the publicly available chronic in vivo calcium imaging dataset of neuronal populations in the auditory cortex of passively listening mice [38]. This dataset comprises *in-vivo* two-photon calcium imaging recordings from 80 fields of view located in auditory cortex of awake mice (N=10). During the recordings, 34 brief stimuli (50 - 70 ms) were presented to the animals, out of which 19 were pure tones and 15 were complex sounds. Each stimulus was presented 20 times in random order. Single neuron activity was calculated as the mean dF/F_0_ in the 40 ms following stimulus onset. Neural activity was recorded in two-day intervals over the time span of one week. On day four (between imaging session two and three), a fear conditioning procedure was carried out by pairing a mild foot shock with a conditioned stimulus. The conditioned stimulus was a complex sound chosen from the 34 aforementioned stimuli. Prior to the recordings, mice were habituated to head fixation and pre-exposed to the set of sound stimuli for at least 5 days to ensure that adaption to novel sensory responses was largely completed and that data acquisition occurred under behaviorally and environmentally familiar and constant conditions. The pre-processing of the data is described in extensive detail in [5]. The pre-processing pipeline is available online via [56].

### Signal Correlation (Figs. 1d-f,j,k, 2a-c, 3a-f,h,i, 4b-i, S1-S10)

Pairwise signal correlations were calculated by computing the Pearson correlation between two neurons’ tuning curves *Γ*_*i*_ and *Γ* _*j*_:

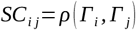

Tuning curves were defined as the stimulus specific median responses over trials.

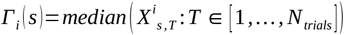

Here, 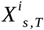 is the average activity over 5 time bins (1 second) after onset of stimulus s in trial T. To account for small sample size effects, we cross-validated the signal correlations using a bootstrapping procedure. To do so, we randomly sampled (with replacement) 20 trials for each stimulus and computed bootstrapped tuning curves for each neuron and signal correlations for each neuron pair. We repeated this N=100 times to obtain N=100 signal correlations for each neuron pair. For all subsequent analyses, we used the mean of these signal correlations.

### Noise Correlation (Figs. 1g-k, 2d-f, 3a-f,h,i, 4b-i, S1-S10)

As a measure of effective recurrent connectivity, we computed pairwise noise correlations by calculating the Pearson correlation between two neurons’ trial-to-trial fluctuation *ζ* _*i*_ and *ζ* _*j*_ over all trials and stimuli:

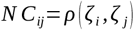

Trial-to-trial fluctuations were computed by subtracting the stimulus specific median response from each single trial response:

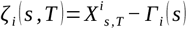

To obtain a more robust estimate of the noise correlation, we applied a bootstrapping procedure similar to the one used for the signal correlation. For each neuron pair, we randomly sampled (with replacement) 20 trials of each stimulus and computed both neuron’s trial-to-trial fluctuation from these samples, using the median obtained from these samples. Bootstrapped noise correlations were then calculated as the Pearson correlation of the bootstrapped trial-to-trial fluctuations. As noise correlations require the trials to be time locked, we used the same trial identities for both neurons when sampling. We then repeated these steps for N=100 times for each neuron pair and used the mean as the neuron pair’s noise correlations.

### Correlation Stability (Figs. 2, S6, S8)

To analyze the stability of pairwise correlations, we first divided neuron pairs into eight groups according to their correlation on a given day. Groups were defined using the following thresholds: (−0.2,0], (0,0.1], (0.1,0.2], (0.2,0.3], (0.3,0.4], (0.4,0.5], (0.5,0.6] and (0.6,1]. Neuron pairs were assigned to a group, when their signal/noise correlation was larger than the lower threshold and smaller or equal to the upper threshold. Correlation stability was then assessed by comparing the mean correlations for these groups between imaging day 1 and 2 (Fig. 2c, f). For correlations less than −0.2, the number of neuron pairs per subgroup (see **Predictive effect of noise correlations on future signal correlations**) was very small (<10 for some subgroups) and deemed insufficient for further analysis (see Fig S3).

### Predictive effect of signal correlations on future noise correlations (Fig. 3b,c, S2)

To assess the predictive effect of signal correlations on noise correlations, we compared the noise correlations on imaging day t+1 of neuron pairs with different signal correlation on day t. For this, neuron pairs were first grouped by their noise correlation on a given day t. In each group, neuron pairs were then further subdivided into subgroups based on their signal correlation on day t. We then computed each subgroup’s mean noise correlation on day t, as well as day t+1 and subsequently compared the noise correlations on day t+1 among the subgroups with the same noise correlation on day t.

### Predictive effect of noise correlations on future signal correlations (Fig. 3e,f, S2)

To assess the predictive effect of noise correlation on signal correlations, we used the same approach as above, but first grouped neuron pairs based on their signal correlations on imaging day t, then subgrouped them by their noise correlation on day t. We then compared the signal correlations on day t+1 of subgroups with the same signal correlation, but different noise correlations on day t.

### Prediction Index (Figs. 3g-i, 4b-i, S6, S8, S9)

The prediction index measures the predictive effect of a predictor variable *X* on a target variable *Y* in an ensemble with *N* elements, where each element *i* has both an *X*-value *X*_*i*_ and a *Y*-value *Y* _*i*_. To calculate the prediction index, we first assigned each of the *N* elements to one of *K* groups by binning them by their *X*-value. An element *i* was assigned to group *k*, if its *X*-value *X*_*i*_ falls into the group specific interval [*θ*_*k, low*_, *θ*_*k, up*_), where *θ*_*k, low*_ denotes the lower and *θ*_*k, up*_ the upper threshold of group *k*.

We then calculated the mean *Y*-value *C*_*k*_ for each of the *K* groups

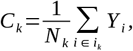

where *N*_*k*_ is the number of elements in group *k* and *i*_*k*_ denotes the index of the elements in group *k*. The prediction index is then the sum over differences between the means of neighbouring groups:

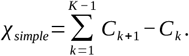

We used the prediction index to compute the influence of signal correlations at one time point on noise correlations at the next time point and vice versa. To do so, we defined the groups on time point t and calculated the group averages at time point t+1:

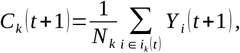

where *i*_*k*_ (*t*) signifies that the groups were defined at time point t, while *Y* _*i*_ (*t* +1) signifies that the values are taken at time point t+1. In addition, we needed to account for a possible a priori spread. For this, we calculated the group averages on time point *t*

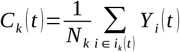

and calculated the difference between the spread on time point t and t+1:

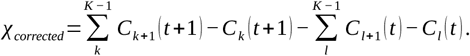

To obtain shuffled prediction indices, we randomly shuffled the group assignment of the neuron pairs.

### Conditional Correlation Analysis (Figs. S4, S5, S7, S10)

To quantify the dependency of noise correlation stability on signal correlations we first grouped neuron pairs by signal correlation on day t, using the same thresholds as before (see section Correlation Stability). For each group, we then calculated the noise correlation stability as the Spearman rank correlation between the noise correlations on day t and noise correlations on day t+1. The groups’ Spearman rank correlation values were then fitted against their mean signal correlations on day t using a linear fit:

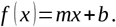

Here, *x* is each group’s mean signal correlation on day t and *f* (*x*) is each group’s Spearman rank correlation between their noise correlation on day t and t+1. To quantify the dependency of signal correlation stability on noise correlation on day t, we proceeded analogously, but with reversed contingencies. Specifically, we grouped by noise correlation on day t and calculated the signal correlation stability between day t and day t+1. In this case, *x* is each group’s mean noise correlation on day t and *f* (*x*) is each group’s Spearman rank correlation between their signal correlation on day t and t+1. From this, the slope *m* was extracted and plotted for different days.

In the model, we fitted the noise (signal) correlation stability to the absolute value of the signal (noise) correlation at time t:

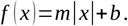

This was done due to the fact that the Hopfield-like mechanism leads to symmetric amplification of positive and negative activities alike. Hence, in the model, the noise (signal) correlation stabilities also increased for increasingly negative signal (noise) correlations, in contrast to the experimental findings.

### Network model (Figs. 4, S8, S9, S10)

Populations of cortical neurons were modelled using a linear recurrent network consisting of N=250 rate neurons with random recurrent connections receiving external input (see Fig. 4a, S8a – S8c). The firing rate dynamics obeyed the differential equation

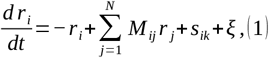

where *r*_*i*_ is the firing rate of neuron *i, M*_*ij*_ describes the recurrent weight from neuron *j* to neuron *I, s*_*ik*_ is the time-independent external input from stimulus *k* to neuron *i*, and *ξ* denotes Gaussian noise independent for each trial, neuron, and stimulus. The simulation was run with 34 different stimuli, which were presented each for 20 trials, analogously to the experiment. The entries of the *N*_*neurons*_ *× N*_*stim*_ stimulus matrix *s* were drawn from a univariate Gaussian distribution with 0 mean and SD=1 (see Fig. S8a for an example). Network weights *M* were normalized such that the maximum Eigenvalue *λ*_*max*_ equals 0.85. This was done by dividing 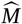 by its maximum Eigenvalue and subsequently multiplying by 0.85.

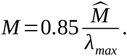

Here, 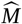 denotes the network weight matrix before normalization. For all model analysis, we used the steady state solution of equation (1), which is given by

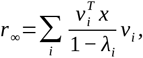

where *x*=*s*_*k*_ +*ξ* is the input vector (of length *N* _*neurons*_) for a given trial of stimulus *k*, and *v*_*i*_ and *λ*_*i*_ are the *i*-th eigenvector and eigenvalue, respectively, of *M*.

Signal and noise correlation matrices (S8d, S8e) and the prediction index (Fig. 4, S8i-n) were computed in the model analogously to the experimental data, but instead of fixed correlation values as thresholds, percentile thresholds were used, to account for differences in the distributions of signal and noise correlations. We used the following percentile thresholds: (0%,20%], (20%,40%], (40%,60%], (60%,80%], (80%,100%].

### Network initialization for plasticity analyses (Figs. 4, S8)

We simulated three different plasticity scenarios, as detailed in the next sections: (I) stochastic drift, (II) Hebbian-like plasticity, and (III) stochastic drift and Hebbian-like plasticity. In models without Hebbian-like plasticity, the weight matrix *M* was initialized as a symmetric matrix with weights drawn from a univariate Gaussian distribution with 0 mean and SD=1. In the models with a Hebbian-like plasticity mechanism, *M* was initialized as the mean over the outer products of 34 randomly generated stimulus vectors *z*_*k*_ (see Fig. S8b for an example):

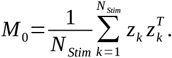

These different initializations were chosen to keep the statistics of the weight matrix similar over the time course of connectivity changes.

### Stochastic drift in the model (Figs. 4c-e,g-i, S8, S9, S10)

Stochastic drift in the input or in the network weights was implemented with an Ornstein-Uhlenbeck (OU) process [41], which allows to introduce stochastic changes in a variable while keeping the distribution stationary over time. The change in the random variable *X* is described as

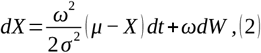

where *μ* and *σ* ^2^ are the mean and the variance of the steady state distribution (and chosen accordingly), *ω* is a free parameter that controls the strength of drift and *W* is a standard Wiener process [57]. To model drift in the input, we applied equation (2) independently to each element of the input vectors *s*_*k*_ varying the strength of drift *ω*. To model drift in the recurrent network matrix *M*, we applied equation (2) to each element *M*_*i, j*_ with *i ≤ j* and then mirrored the values, *M* _*ji*_ :=*M* _*ij*_, for *i*> *j* to yield a symmetric connectivity matrix *M*. Again, we simulated different drift rates by varying *ω*.

### Hebbian-like plasticity in the model (Figs. 4f-i, S8, S9, S10)

Input specific plasticity in the recurrent network weights was implemented via a Hebbian-like mechanism by which network weights were adapted based on the input they receive. Recurrent network weights were strengthened when neuron pairs received correlated input using the following learning rule (inspired by Hopfield [58])

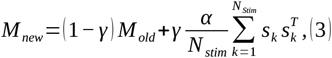

where *M*_*new*_ is the new connectivity matrix, *M*_*old*_ is the previous connectivity matrix, and *γ* is the learning rate. The constant *α* was chosen such, that the standard deviation of 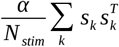 was equal to that of *M*_*old*_. After each update step, *M*_*new*_ was normalized such that its maximum eigenvalue was equal 0.85. This mechanism has one free parameter, the learning rate *γ*. We varied *γ* to control the speed at which new stimuli are incorporated into the network weights.

### Models with Hebbian-like plasticity and stochastic drift (Figs. 4h-i, S8, S9, S10)

In models with both Hebbian-like plasticity and stochastic drift in the recurrent connections, the network weights were adapted by adding a matrix *dΩ* to the right-hand side of equation (3):

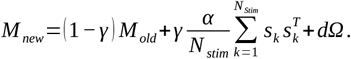

The elements of *dΩ* represent changes due to an OU-process and were computed at each update step as

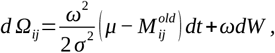

for each pair *i, j* : *i ≤ j* and then mirroring the values, *Ω*_*ji*_ :=*Ω*_*ij*_, for *i*> *j* to yield a symmetric matrix*M* _*new*_.

### Model statistics (Figs. 4, S8, S9, S10)

With any of the above mechanisms, the simulation was run for *N*_*t*_ =100 update steps, one update step being defined by the iterative equations (2) and (3) in the previous sections. One update step models the transition from one imaging day to the next in the experiments. We systematically varied the free parameters in the scenarios described above (*γ, ω*, or both, respectively) and repeated this procedure *N* _*init*_ =10 times for each parameter setting. The mean prediction indices shown in (Fig. 4c-4i, Fig. S9) represent means over all signal or noise correlation groups, update steps and initializations. The slopes shown in Fig. S10 represent the means over all update steps and initializations.

### Quantification and statistical analysis

All statistical analyses were performed using MATLAB (Mathworks, Natick, MA, USA). We used the following statistical tests for the given statistical analyses.

### Comparison between prediction indices (Figs. 3h, 3i, S6)

To compare the prediction indices from different days or contingencies, the prediction indices were tested for significance using a two-sided Wilcoxon signed rank test. The p-values were subsequently corrected for multiple comparisons with a Bonferroni correction. Star denotes significance *p<0.05.

### Comparison between parameter fits (Figs. S4, S7)

To test for significant differences between fitted parameters across days, a distribution of parameter values was generated using a bootstrapping procedure. N=100 surrogate datasets were generated by randomly sampling neuron pairs with replacement from the original dataset. Next, the conditional correlation analysis was performed on each of the surrogate datasets resulting in a distribution of slope values *m* for each day and contingency. Significance was tested using a two-sided Wilcoxon-Mann-Whitney test. The p-values were subsequently corrected for multiple comparisons with a Bonferroni correction. Star denotes significance *p<0.05.

## Author Contributions

Conceptualization: MK, JBE; data curation: DA; formal analysis: TL; funding acquisition: SR, MK; investigation: TL, JBE; methodology: JBE, TL; project administration: MK; resources: MK, SR; software: TL; supervision: MK, SR, JBE; validation: TL, JBE; visualization: TL; writing - original draft: JBE, TL; writing - review & editing: SR, MK.

## Acknowledgements

We thank members of the Kaschube and Rumpel lab for helpful discussions. This work was supported by research grants Deutsche Forschungsgemeinschaft CRC1080-C05, Deutsche Forschungsgemeinschaft SPP 2041 Project #347573108, Deutsche Forschungsgemeinschaft/Agence nationale de la recherche Project #431393205 and Deutsche Forschungsgemeinschaft DIP “Neurobiology of Forgetting”. We want to acknowledge the assistance of ChatGPT (OpenAI) in the revision and refinement of this manuscript.

## Ethical Statements

All animal experiments were performed in accordance with the Austrian laboratory animal law guidelines for animal research and had been approved by the Viennese Magistratsabteilung 58 (Approval M58/00236/2010/6).

## Code and Data Availability

Data and code used for the presented analysis is publicly available via https://gin.g-node.org/thomaslai/RD-SC-NC

