## Supplementary Information for "Representational drift reflects ongoing balancing of stochastic changes by Hebbian learning"

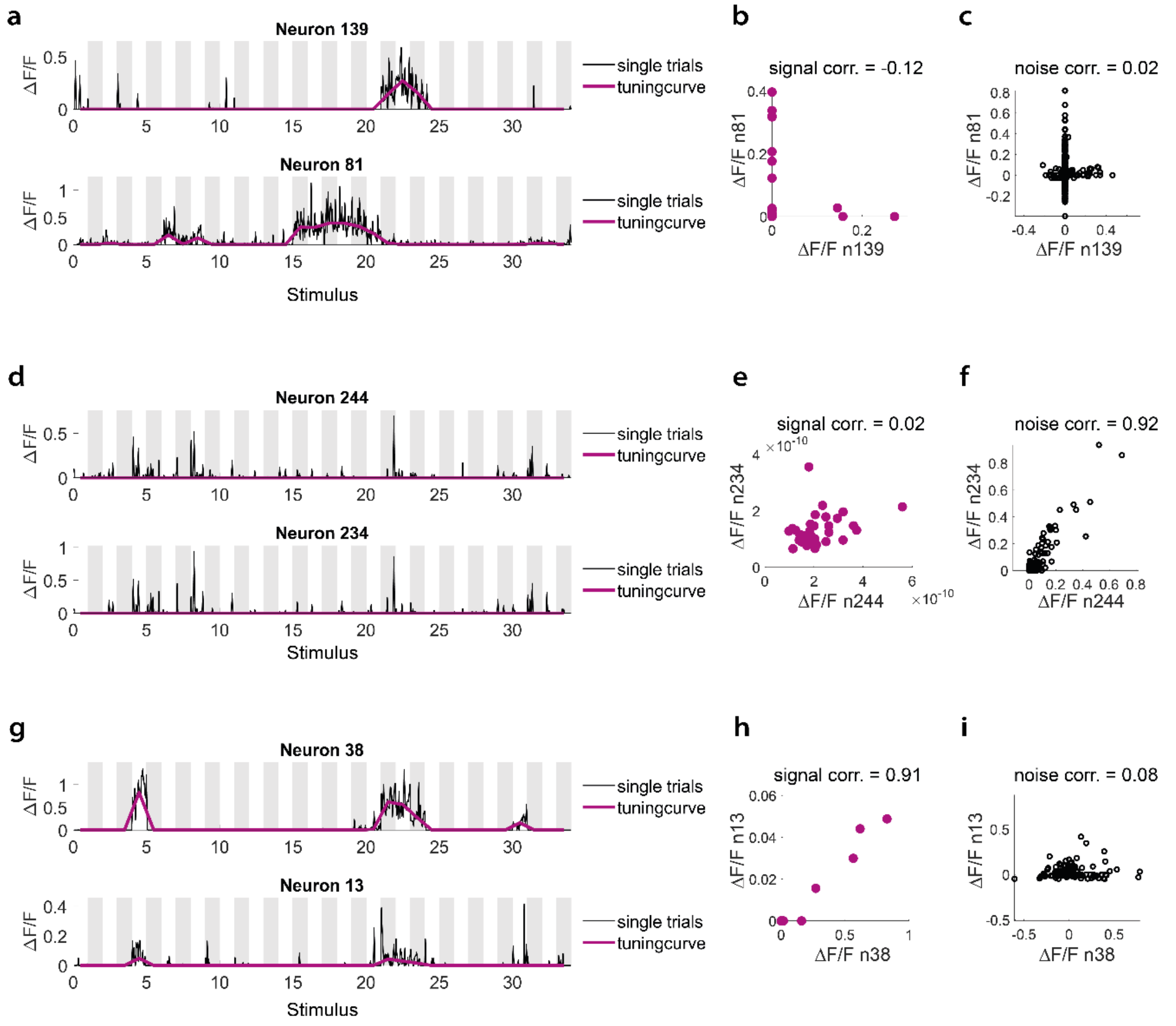


**Figure S1: Example neuron pairs.**
a)-c) Neuron pair with low signal correlation and low noise correlation. a) Single trial response (black thin line) and stimulus specific median response (thick purple line) of both neurons. b) Stimulus response of the first neuron vs. the second neuron. c) Trial-to-trial fluctuation around the median response of the first neuron vs. the second neuron. d) - f) Same as a) - c), but for a neuron pair with low signal correlation and high noise correlation. g) - i) Same as a) - c), but for a neuron pair with high signal correlation and low noise correlation.

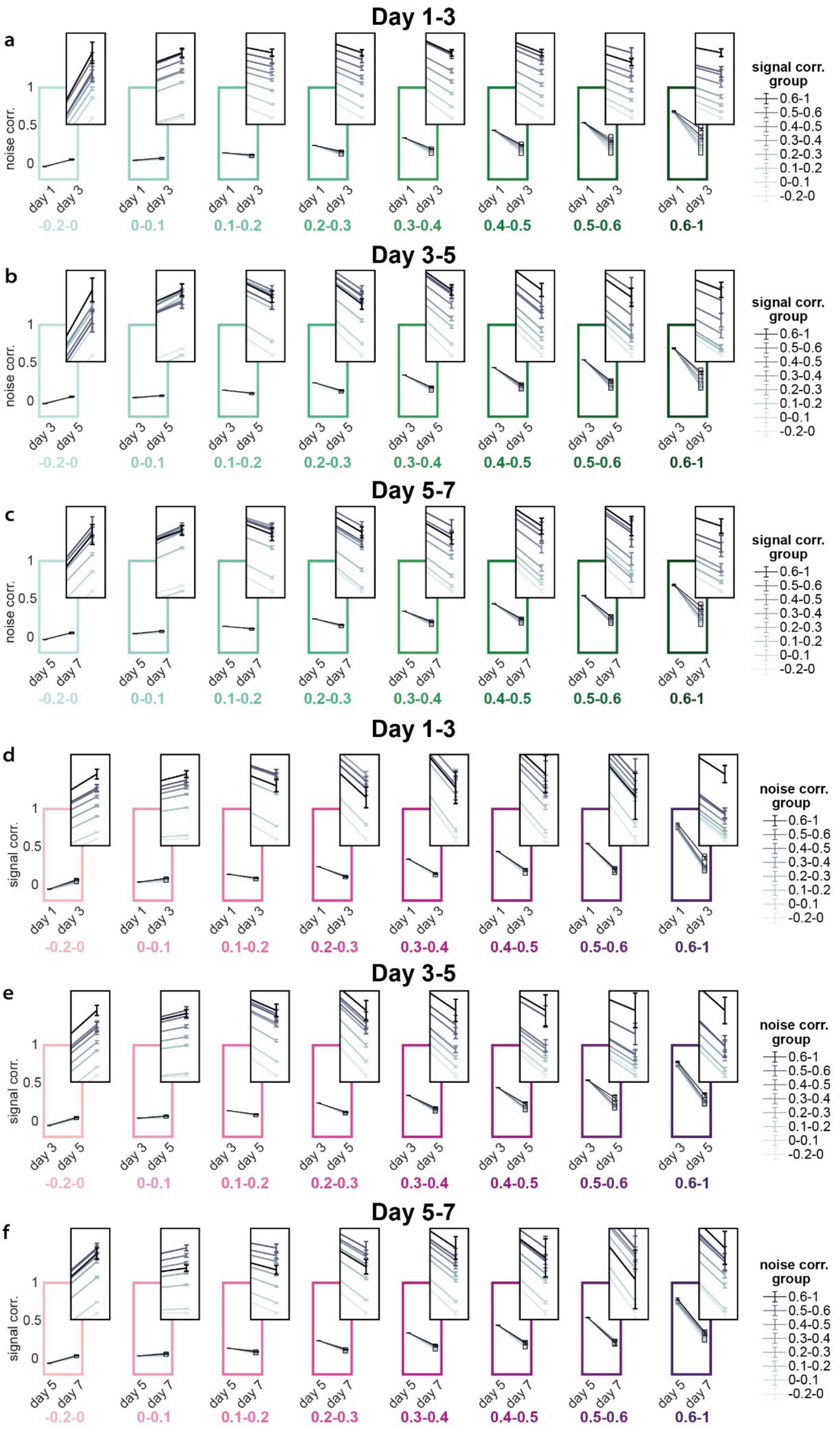
Day 1-3
a
noise corr.
1
0.5
0
day 1
day 3
-0.2-0
0-0.1
0.1-0.2
0.2-0.3
0.3-0.4
0.4-0.5
0.5-0.6
0.6-1
signal corr. group
0.6-1
0.5-0.6
0.4-0.5
0.3-0.4
0.2-0.3
0.1-0.2
0-0.1
-0.2-0
Day 3-5
b
day 3
day 5
Day 5-7
c
day 5
day 7
d
signal corr.
noise corr. group
e
f

**Figure S2: Predictive effect of signal correlation on noise correlation, vice versa, for all groups and days.**

a) – c) Mean noise correlation of neuron pairs grouped by their noise correlation (individual columns represent individual signal correlation groups) and subgrouped by their signal correlation (individual lines represent individual subgroups) on day 1-3 (a), day 3-5 (b) and day 5-7 (c). d) – f) Mean signal correlation of neuron pairs grouped by their signal correlation (individual columns represent individual noise correlation groups) and subgrouped by their noise correlation (individual lines represent individual subgroups) on day 1-3 (d), day 3-5 (e) and day 5-7 (f). Error bars represent SEM.

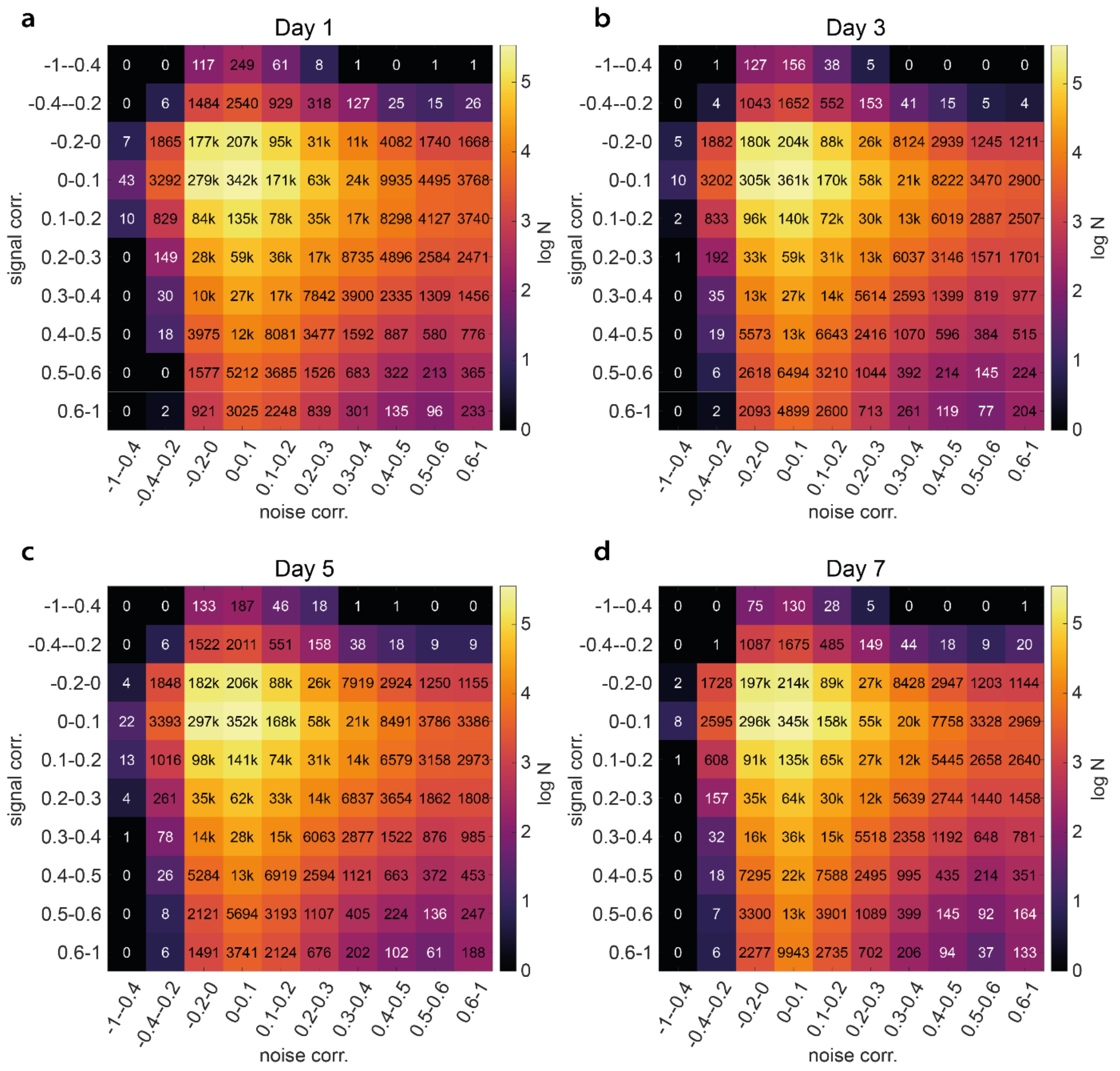


**Figure S3: Distribution of neuron pairs across the subgroups.**

Number of neuron pairs in each subgroup on day 1 (a), day 3 (b), day 5 (c) and day 7 (d).

**Signal correlations stabilize noise correlations, but not vice versa**

In the Results section we showed that high signal correlations on day 1 predicted a reduced decay in noise correlations from day 1 to day 3, indicating a predictive effect of signal correlations on future network connectivity. However, prediction alone does not fully capture the dynamic relationship between signal and noise correlations. Here, we introduce a complementary measure that focuses on stabilization rather than prediction, allowing us to assess how signal correlations contribute to the maintenance of network connectivity over time.

To quantify stabilization more directly, we again grouped neuron pairs by their signal correlations on day 1. For each of these groups we then assessed noise correlation stability by calculating Spearman's rank correlation between noise correlations on day 3 and noise correlations on day 1 (Fig. S4a, for all groups see Fig, S5a). This analysis revealed an almost linear relationship between signal correlation and noise correlation stability (Fig. S4b), suggesting that signal correlations contribute to the retention of effective connectivity between neuron pairs. To quantify this effect, we fitted noise correlation stability to signal correlations using a linear fit $f(x) = mx + c$. The slope $m$ of this linear fit is shown in Fig. S4c. When neuron pairs were shuffled across groups with similar signal correlations, this dependency disappeared (Fig. S4b, S4c).

Conversely, when we grouped neuron pairs based on their noise correlations and scattered their signal correlations on day 3 against those on day 1, no stabilizing effect was observed. Although signal correlation stability was generally higher than noise correlation stability, it showed only a weak negative linear dependence on the noise correlation group (Fig. S4d-f, for all groups see Fig. S5d). The overall increased stability and lack of linear dependence were also evident in the Spearman's rank correlation (Fig. S4e) and in the slope $m$ of the linear fit $f(x) = mx +c$ (Fig. S4f).

Together, these results indicate that neuron pairs with high signal correlation tend to exhibit more stable noise correlations than pairs with low signal correlation, In contrast, the stability of signal correlation appears to be only minimally influenced by the level of noise correlation.

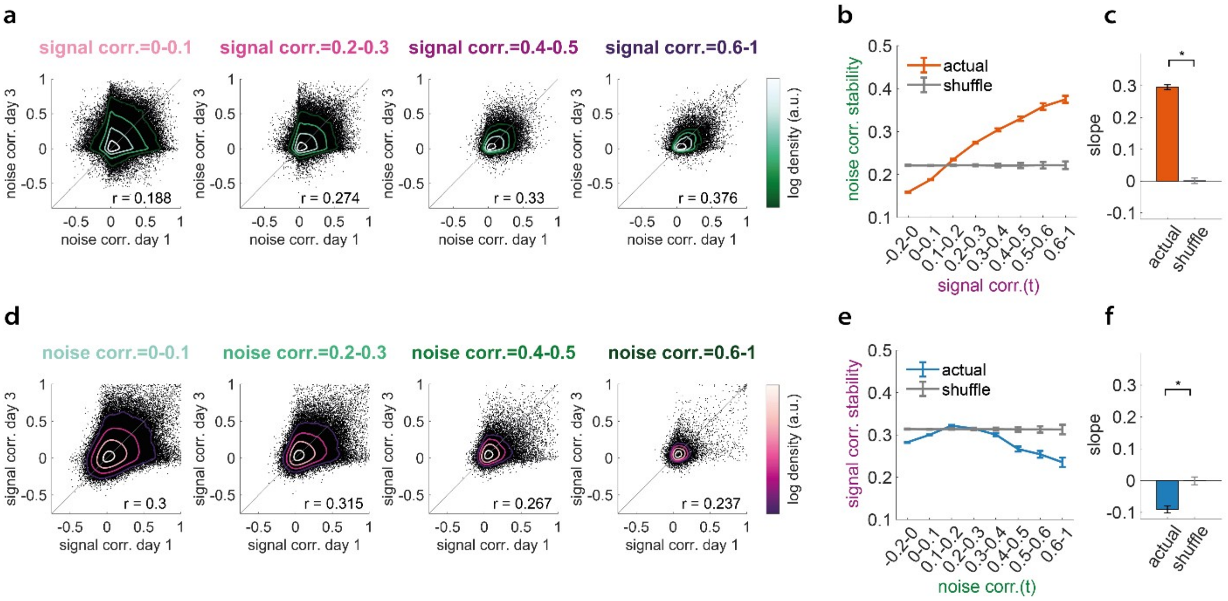


**Figure S4: Signal correlation stabilizes noise correlation under basal conditions.**
a) Noise correlation on day 1 vs. noise correlation on day 3 for groups of neuron pairs with different signal correlation on day 1. Shown are 4 out of 8 groups. b) Noise correlation stability measured as the Spearman rank correlation between noise correlation on day 1 and day 3 for each signal correlation group (mean +- SD of bootstrapped distribution). Shuffle in gray. c) Slope obtained from linear fit of noise correlation stability to signal correlation on day t. d)-f) Same as (a)-(c), but with switched signal and noise correlations. *p<0.05 (MWU-test).

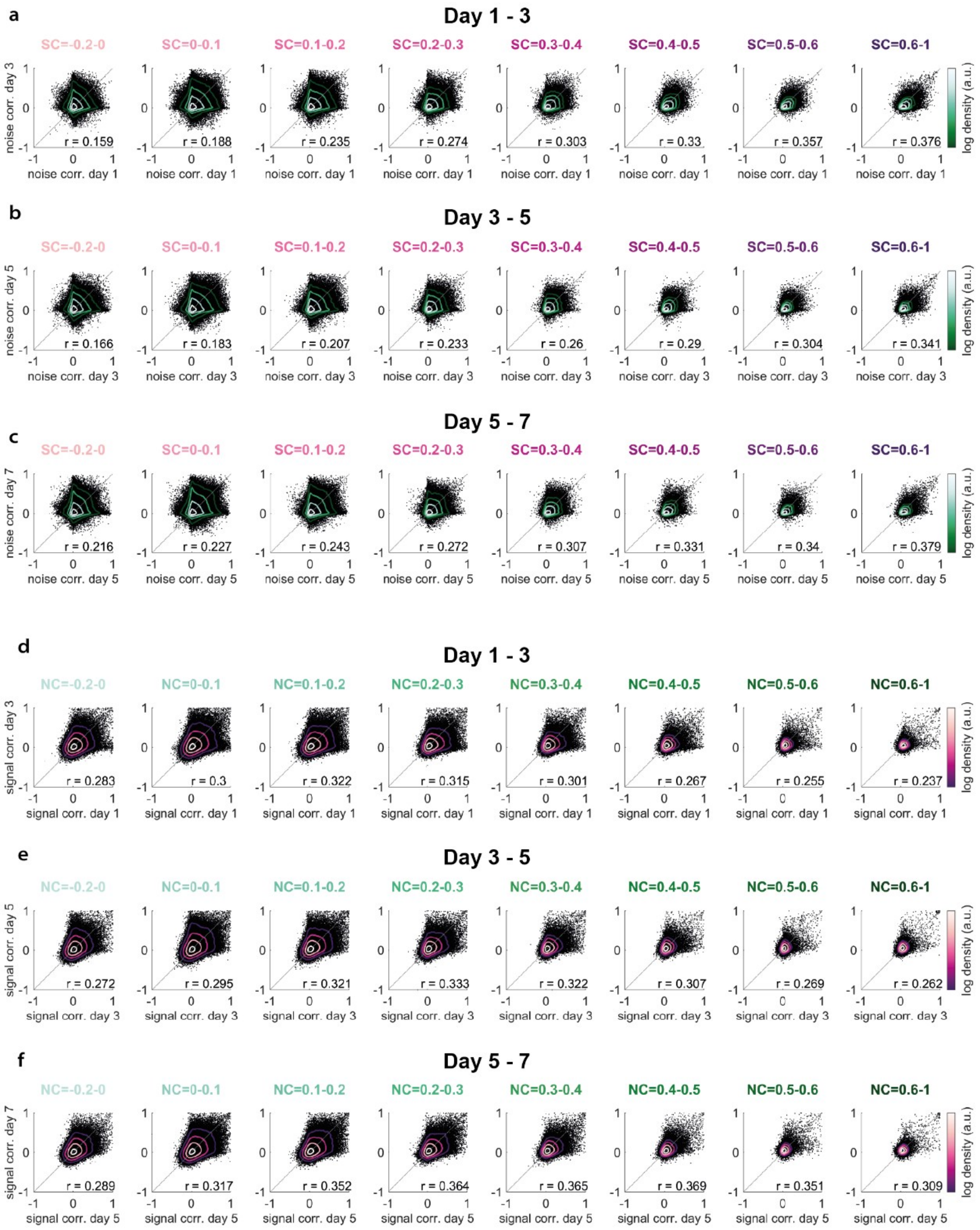


**Figure S5: Conditional signal - and noise correlation stability on all days.** Noise correlation on day t vs. noise correlation on day t+2, grouped by signal correlation on day t. a) Day 1 – 3, b) day 3 – 5, c) day 5 – 7. Signal correlation on day t vs. signal correlation on day t+2, grouped by noise correlation on day t. d) Day 1 – 3, e) day 3 – 5, f) day 5 – 7.

**The impact of fear conditioning on signal and noise correlations**

In the Results part, we have examined behaviourally stable conditions and found evidence of ongoing Hebbian-like learning. But what happens during actual behavioural learning? To address this, we investigated the dynamics of signal and noise correlations during auditory cued fear conditioning (ACFC), a well-established paradigm that induces behavioral learning. This allowed us to assess how this type of learning directly affects these correlations. ACFC has been shown to impact synaptic connection dynamics in the auditory cortex [32], as well as the dynamics of sound-evoked activity patterns [5], [32], [35], [36], [37]. Given that Hebbian-like plasticity is often associated with behavioral learning, we expected to see an increase in both the predictive power and stabilizing effect of signal correlations on noise correlations, a connection which has also been supported by a recent modeling study [59].

To test this, we applied auditory cued fear conditioning between the second and third imaging session by pairing one of the stimuli with a mild electric shock. The animals successfully learned this association, as demonstrated by their freezing behaviour when presented with the conditioned sound in a neutral context (for details see [5]). When computing signal and noise correlation stability as before, we found no significant effect of fear conditioning on the volatility of signal and noise correlations. Similarly, the prediction indices were not significantly affected in either direction (Fig. S6). However, when computing conditional correlations, we observed that the stability of noise correlations continued to exhibit a linear relationship with signal correlations, but this dependency weakened during and after ACFC. Specifically, the slope of the linear fit of noise correlation stability to signal correlation showed a significant reduction (Fig. S7a, S7b). This indicates that, while signal correlation remained a predictor of noise correlation stability, its influence diminished after ACFC. Conversely, when analysing how noise correlations affect the stability of signal correlations, an effect that was initially weak under basal conditions, we observed a significant amplification following fear conditioning (Fig. S7c, S7d): Both the overall stability (indicated by the y-axis intercept) and the dependency of signal correlation stability on noise correlations (reflected by the slope) were progressively enhanced after ACFC. This suggests that the influence of noise correlations on the stability of signal correlations became more pronounced after fear conditioning. In the model this was achieved by a reduced learning rate compared to stochastic drift rates.

Thus, counterintuitively, rather than seeing an increase in Hebbian-like learning during fear conditioning, we observed a reduction in its effect on noise correlations. This may suggest that less ongoing learning occurs during behavioural learning to facilitate the consolidation of learned associations into memory storage.

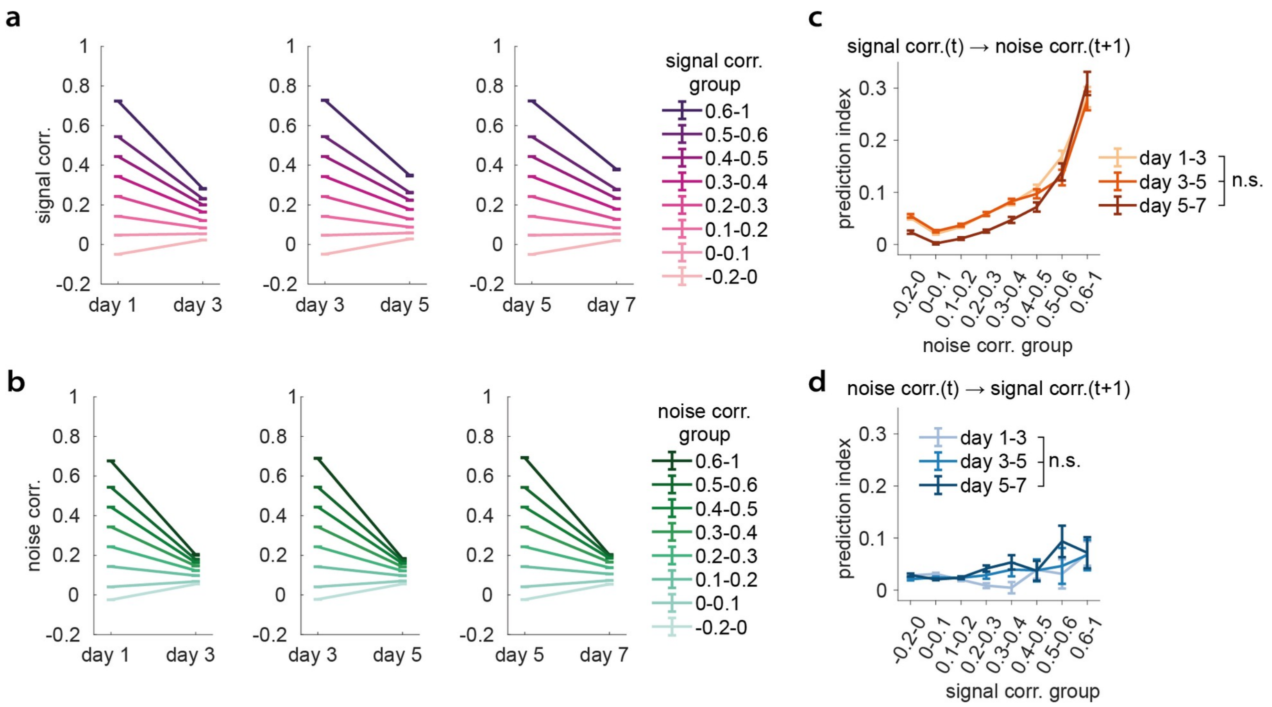


**Figure S6: Fear conditioning has little effect on the stability and predictive effect of signal and noise correlations.** a) Signal correlation stability under basal conditions (left) and during fear conditioning (middle and right). b) Noise correlation stability under basal conditions (left) and during fear conditioning (middle and right). Error bars represent SEM. c) Prediction index of the noise correlation based on signal correlations. d) Prediction index of the signal correlation based on noise.

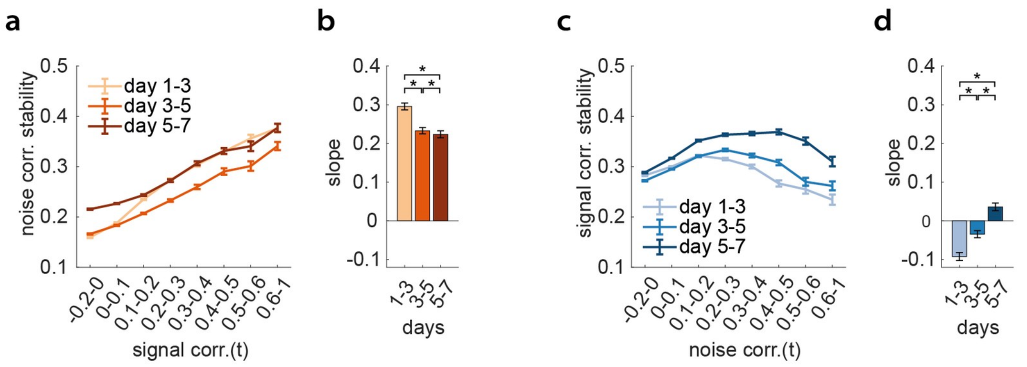


**Figure S7: Fear conditioning reduces the predictive effective effect of signal correlations on future noise correlations.**
a) Spearman rank correlation between noise correlation on imaging days t and t+1 w.r.t. the signal correlation on day t. b) Slope from linear fit of noise correlation stability to signal correlation on day t. c) Spearman rank correlation between signal correlation on day t and day t+1 w.r.t. the noise correlation on day t. d) Slope from linear fit of signal correlation stability to noise correlation on day t. Error bars represent SD of bootstrapped distribution. n.s. non-significant, *p<0.05 (MWU-test).

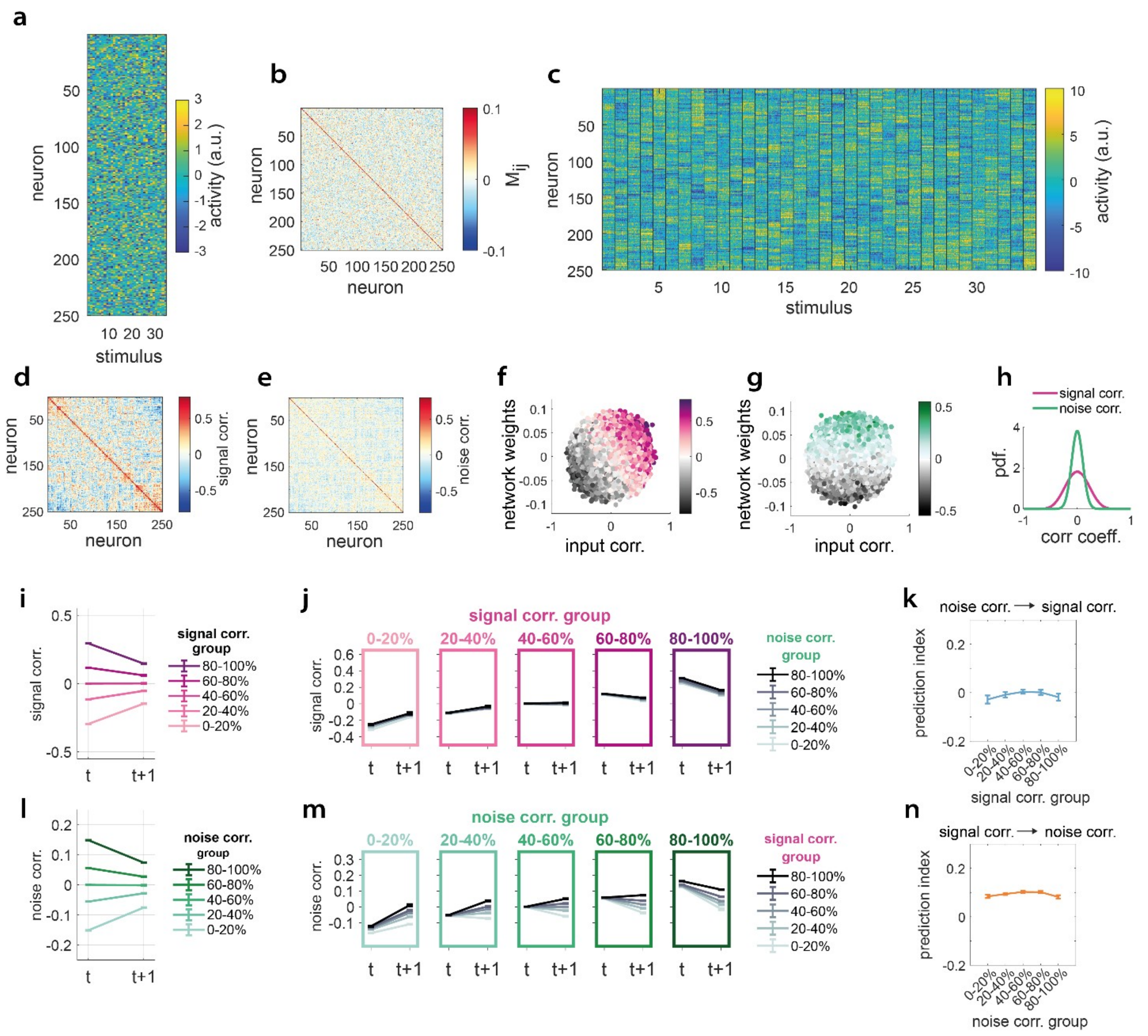


**Figure S8: Example data of a model with input drift ($\omega_{\square}$= 1.1), network drift ($\omega_{net}$= 0.4) and network plasticity ($\gamma$=0.16).**

Data from timepoint $t=25$ of the simulation is shown unless stated otherwise. a) Example of external input into the network. b) Example of recurrent network weights. c) Output activity for the input and network given in a) and b). Black lines indicate different stimuli, individual columns indicate single trials. Signal correlation matrix (d) and noise correlation matrix (e) obtained from the activity shown in c). f) and g) Network weights vs. input correlation, pooled over all days. Colour coded is the signal correlation (f) and the noise correlation (g). h) Distribution of signal correlations and noise correlations pooled over all timepoints. i) Mean signal correlation of neuron pairs sorted into groups by their signal correlation. Errorbars represent SEM. j) Mean signal correlation of each signal correlation group (individual columns) subgrouped by their noise correlation (individual lines). k) Prediction index of noise correlations for signal correlations. Error bars represent SD of bootstrapped distribution. l) - n) Same as i) - k) but for the predictive effect of signal correlations on noise correlations.

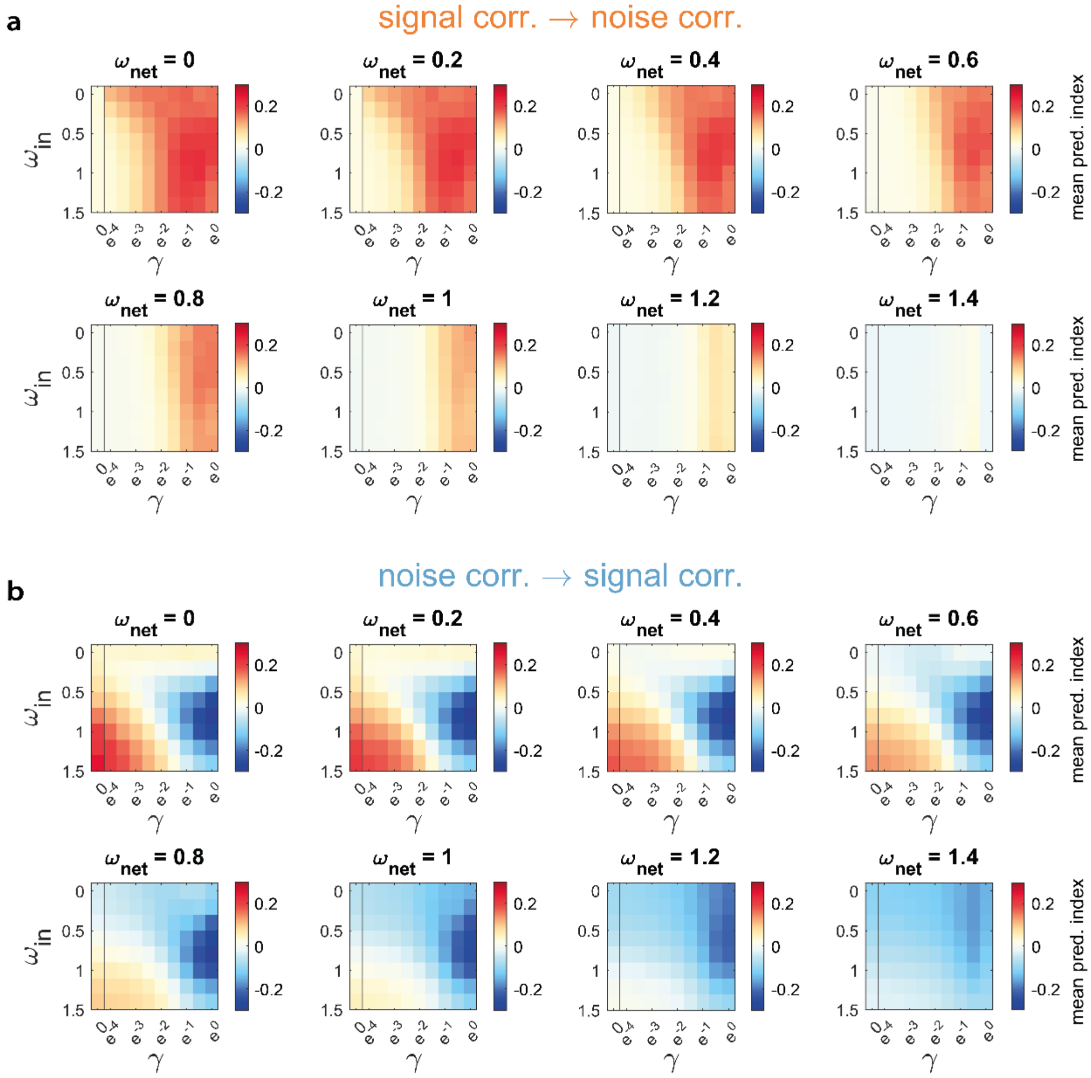


**Figure S9: Parameter scan of prediction index in model with input drift, network drift and network plasticity.**

a) Mean prediction index of signal correlations on noise correlations. Individual subplots show different values for network drift $\omega_{net}$. b) Mean prediction index of noise correlations on signal correlations.

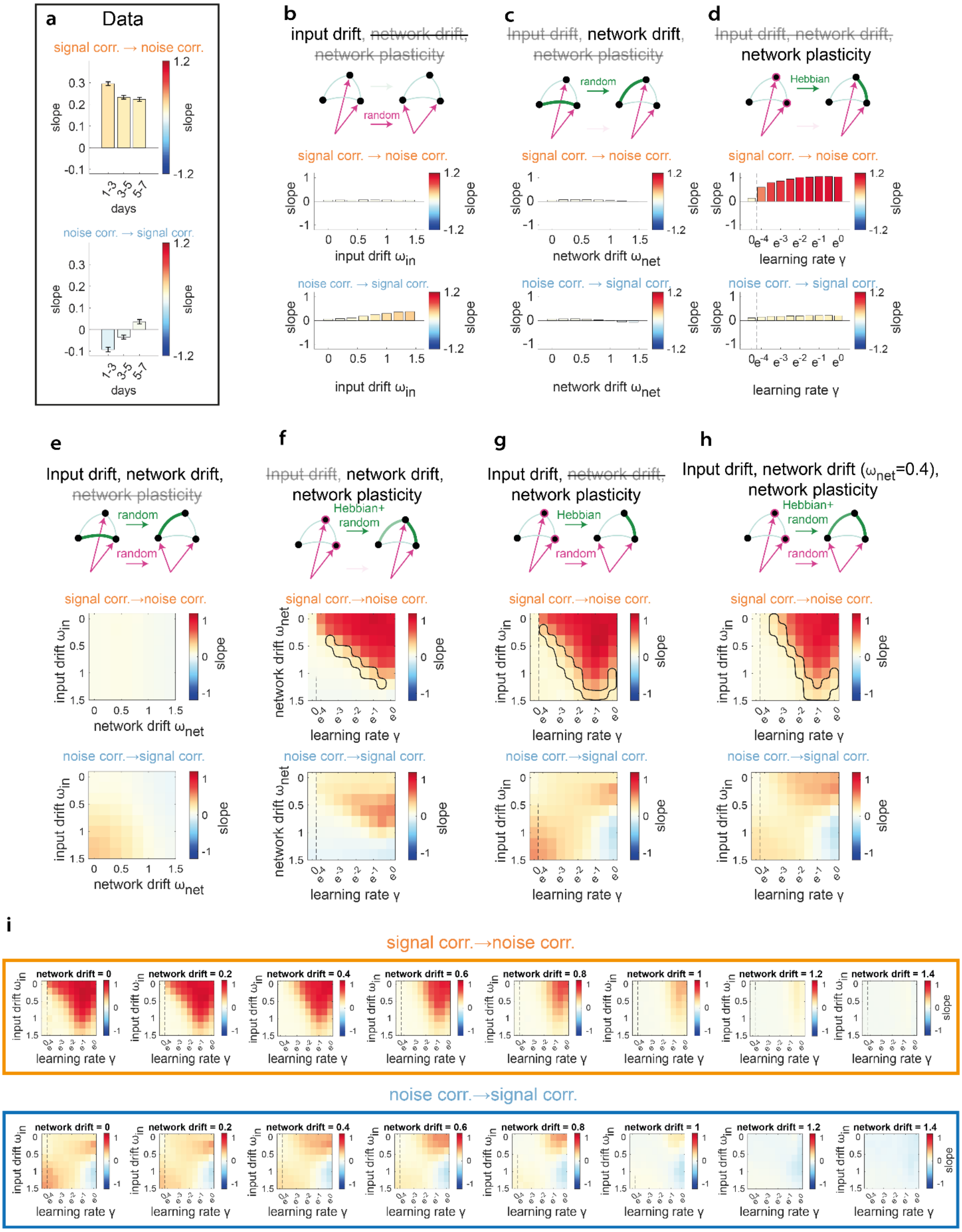


**Figure S10: Parameter scan for conditional correlation stability analysis in model.**

Slope value obtained from linear fit of signal (noise) correlation stability from time t to t+1 to noise (signal) correlation at time t. a) Slope obtained from the experimental data over days. Top: Noise correlation stability w.r.t. signal correlation at time t; bottom: Signal correlation stability w.r.t. noise correlation at time t. Errorbars represent SD of bootstrapped distribution. b) – d): Parameter scan of models with one mechanism. b) Model with only input drift. c) Model with only network drift. d) Model with only network plasticity. e) – g): Parameter scan of models with two mechanisms. e) Model with input drift and network drift. f) Model with network drift and network plasticity. g) Model with input drift and network plasticity. h) – i) Parameter scan of model with all three mechanisms. h) Parameter scan in the input drift – learning rate plane for $\omega_{net}=0.4$. i) Parameter scan for all $\omega_{net}$. Top row: slope for noise correlation stability w.r.t. signal correlation at time t, bottom row: slope for signal correlation stability w.r.t. noise correlation at time t. Individual matrices represent different network drift strengths.
